# Global functional connectivity of cognitive control network predicts task-switching performance in older adults

**DOI:** 10.1101/2023.11.20.567605

**Authors:** B. Madero, M. Sodoma, C. Oehler, V. A. Magnotta, J. D. Long, E. Hazeltine, M. W. Voss

**Author notes:** Corresponding author Michelle W. Voss, Ph.D., Associate Professor of Psychological and Brain Sciences, Office: 274 Psychological and Brain Sciences Building, Mail: G60 Psychological and Brain Sciences Building, Iowa City, Iowa 52242-1407.

## Abstract

Executive Function (EF) is a complex higher order cognitive process that involves integrating subprocesses of attention, working memory, inhibition, and task switching and known to be one of the first cognitive functions to be affected by age related declines. Global Functional Connectivity (GFC), a measure of whole brain connectivity in brain networks such as the Cognitive Control Network (CCN), has been shown to measure the maintained EF abilities in older adults with cognitive impairment. However, this relationship has not been explored in a sample comprised of mostly cognitively normal older adults nor with the use of an experimental EF task sensitive to sub-processes most sensitive to aging. Task-switch paradigm studies have shown age-related deficits in holding and responding to rule sets while inhibiting another rule set in working memory (i.e., mixing cost). The present study aims to elucidate the relationship seen in EF performance using GFC and leveraging a task-switch paradigm that manipulates response overlap to increase cognitive demands on switching between tasks (e.g., mixing cost). Results show that greater negative DMN GFC coincides with older adults’ lower mixing costs, not CCN GFC. Findings indicate that DMN GFC, not CCN, may be important for EF performance in older adults.

## 1. Introduction

The ability to manage multiple tasks affects an individual’s capacity to maintain a healthy independent life in aging and avoid accidents during activities that require multi-tasking such as driving and cooking. Executive function (EF), within the broader concept of cognitive control, reflects the individual’s ability to efficiently attend to goals, inhibit actions inconsistent with our goals, and plan future tasks (Badre, 2008; Mackie et al., 2013). EF abilities decline and begin to vary with some individuals having poorer performance than others (Tucker-Drob, 2019). Age-related EF declines co-occur with frontal cortex structural atrophy and weaker connectivity of brain networks implicated in EF processes (Lacreuse et al., 2020), though the exact mechanism between these relationships is unknown. With an aging population continuing to grow, it is imperative to not only understand the mechanistic aging declines of EF but also ascertain the variability in older adults.

The brain is organized into functionally connected brain networks comprised of distinct brain regions that coactivate to support task performance. Research has shown that not only are these networks important for carrying out specific functions but their coactivation strength or functional strength, declines with age (Chan et al., 2014).

One such network, the Cognitive Control Network (CCN), is comprised of dorsolateral prefrontal cortex, anterior cingulate cortex/pre-supplementary motor area, dorsal premotor cortex, anterior insular cortex, inferior frontal junction, and posterior parietal cortex (Cole & Schneider, 2007). CCN brain regions communicate locally within the CCN network and also with other networks distributed throughout the cortex to perform complex tasks known to require memory for hierarchical rule sets and sensorimotor control (Badre, 2008; Chan et al., 2014; Cole & Schneider, 2007). Thus, the CCN communicates with other networks (e.g., ventral attention, sensory motor) to carry out cognitive control functions for goal directed behaviors (Menon & D’Esposito, 2022). The Default Mode Network (DMN) is known to be negatively correlated with the CCN (i.e., disengages as the CCN engages) during performance of a task as it is most engaged when attention is directed inward during memories, imagination, mind wandering, planning, or thoughts about the self (Andrews-Hanna et al., 2014; Sridharan et al., 2008). The negative correlation (i.e., anticorrelation) strength of the DMN with CCN is related to better performance in working memory, memory, and cognitive control (Breukelaar et al., 2020; Kelly et al., 2008; Zou et al., 2013). The sensory motor network is used to enact the action of the planned goal (DMN) and external goal behavior (CCN) and also processes external stimuli information such as finger tapping (Biswal et al., 1995).

It should be emphasized that in order to perform higher order processes of EF, coordination of the CCN must be functionally interconnected with other networks in order to orchestrate desired behaviors efficiently (Friedman & Robbins, 2022). This relationship has been found to not only be with task activation but also with resting-state measures as it has been shown to be the backbone of the functional connectome (Breukelaar et al., 2020; Cole et al., 2014). Global functional connectivity (GFC), also referred to as global brain connectivity or weighted degree centrality (Cole et al., 2011; Cole et al., 2010; Cole et al., 2012; Rubinov & Sporns, 2011), is a measure that accounts for a network’s full-range (i.e., whole cortical brain) of connectivity, within known regions of a network or system and beyond its functional boundaries. It has been shown in healthy young and early middle-aged adults (Cole et al., 2012) that greater CCN GFC is related to better cognitive performance as well as greater maintained EF in mild cognitively impaired (MCI) adults (Franzmeier et al., 2017). However, it is unknown in this line of work how GFC measurement interacts with EF processes where both age-related behavioral declines and age-related functional declines of network strength are observed (Dickerson et al., 2008; Grambaite et al., 2011; Voss et al., 2016).

To measure individual differences in EF and cognitive control broadly, we use a task-switch paradigm designed to engage core cognitive control processes of the CCN. Switching between tasks requires individuals to attend to more than one task, inhibit actions associated with previously irrelevant tasks, hold information associated with multiple different tasks in working memory, and bind external goals to the actions needed to carry out these goals. The paradigm consists of rule-sets that are a combination of visual cues (i.e., color patches) and targets (i.e., digits and letters) mapped to button press responses. Thus, maintenance of multiple rules and their response mappings is essential for successful performance (Mayr, 2001; Rogers & Monsell, 1995).

This paradigm produces two measures related to cognitive control. The cognitive costs of maintaining multiple tasks and switching between tasks can be disentangled by measuring mixing cost (also known as global switch cost) and switching cost (local switch cost). Mixing cost is operationalized as the difference in reaction time between responses in blocks that contain one rule set only (i.e., a single task block) and responses that repeat the rule set from the previous trial within blocks that contain at least two distinct rule sets. Thus, the mixing cost indicates the cost of maintaining multiple rule sets when one or more sets are irrelevant for the current trial. In comparison, switching cost is operationalized as the difference between trials when the rule set repeats from one trial to the next within a block containing multiple rule sets and when the rule set switches, necessarily within a block containing multiple rule sets (Eich et al., 2016; Mayr, 2001; Rogers & Monsell, 1995; Wasylyshyn et al., 2011).

The two types of costs, mixing cost and switching cost, are differentially affected by the relationship between the stimulus-response mappings for the tasks. Seminal findings in Mayr (2001) found that older adults had increased mixing cost when rule set responses overlapped compared when the two rules set responses were separate. That is, when two tasks use overlapping response keys either overlapping stimulus sets (e.g., using the same stimuli for different rules), mixing costs increased in older adults compared to young adults but switch costs are similarly affected in both age groups (Mayr, 2001; Verhaeghen, 2011; Wasylyshyn et al., 2011). However, it is unclear, what brain mechanisms lead to greater mixing cost in older adults when aspects of either/both task rules and responses have high overlap.

The current study aimed to leverage the seminal findings by Mayr (2001) to understand network mechanisms of age-related declines in cognitive control capacity as measured by task-switching processes. We used two experimental conditions that differed in terms of whether the two task sets involved separate or overlapping responses. We predicted that older adults would have larger mixing cost in the Overlap condition compared to the Separate condition by comparing performance within-subjects on Separate and Overlap conditions. What remains unknown, however, is whether differences in resting-state global functional connectivity relates to mixing cost costs in the Overlap condition (high EF demand) compared to the separate (low EF demand) condition. We predicted that CCN GFC measures cognitive control capacity in cognitive normal older adults, greater CCN GFC would be correlated with lower mixing cost compared to switching cost, and this relationship would be magnified under the highest cognitive control demand in the Overlap condition.

## 2. Materials and Methods

### Participants

Participants were recruited as part of a larger intervention trial (NCT03114150), and the methods and data reported here are from the baseline MRI and neuropsychological assessments only. Eligible participants met the following criteria: (1) were between ages of 55 and 80 years old, (2) free from psychiatric and neurological illness and no history of stroke or chronic metabolic diseases, (3) no history of brain trauma or brain surgery, (4) no self-reported regular use of medications affecting the central nervous system (psychotropics, chemotherapy, etc.), (5) >20 on MoCA, (6) able to complete an MRI scan (i.e. no implants above the waist, can comfortably fit in scanner, etc.), (7) corrected vision of 20/40, (8) able to complete a supervised maximal graded exercise test for the exercise intervention portion of the study (not considered here), (9) do not require caregiver assistance, and (10) not colorblind. Participants were recruited from the surrounding area with flyers, posters, emails, and informational stands at local events. All participants provided consent to the study procedures approved by the University of Iowa Institutional Review Board (IRB) and were compensated for their participation at the end of the intervention study. The final sample recruited for this study consisted of 122 adults (66% female, 16.2(2.4) years of education). Average MoCA scores additionally indicated participants were cognitively normal on average, with a mean (SD) score of 27.2 (1.97) out of 30 (Nasreddine et al., 2005).

### Task-switching paradigm

Our task-switching paradigm consisted of two experimental conditions, *Separate* and *Overlap*, completed on different days with the order counterbalanced between subjects. Each condition contained 4 pure blocks (repeated single task) and 2 mixed blocks (mixed task). Each condition consisted of a practice block and test phase block (one for each rule set and block type). Each single task block consisted of 24 trials for both practice and test. In the mixed blocks, the practice block consisted of 48 mixed trials while the test block consisted of 160 trials.

In the *Separate* condition, participants performed either the vowel/consonant letter discrimination task or the odd/even number discrimination task. In the vowel/consonant task participants were asked to determine whether the letter stimulus was a vowel or consonant (vowel: A/U; Consonant: C/W). In the odd/even task, participants were asked to determine whether the number stimulus was odd or even (odd:1/3; even: 2/4). Different colors cued which rule set was to be retrieved from memory to respond (green cue: vowel/consonant task; blue cue: odd/even task). A critical aspect of our manipulation for the task, response mappings were shown on the screen with each rule set corresponding to each hand (vowel/consonant: left hand; odd/even: right hand) using the index and middle finger. Participants also received feedback on each trial of the practice blocks. In the second phase of the condition, participants performed the mixed block in which the rules randomly switched between trials without feedback.

In the *Overlap* condition, participants performed either the odd/even number discrimination task or the straight/curved number discrimination task. In the odd/even task participants were asked to determine whether the number stimulus was an odd or even (odd:1/3; even: 2/4).

In the straight/curved task, participants were asked to determine whether the number stimulus was straight or curved (straight:1/4; curved: 2/3). Different colors cued which rule set determined the response which are stored in memory (blue cue: odd/even task; red cue: straight/curved task). Response mappings were shown on the screen with each rule set corresponding to one hand using the index and middle finger to respond to *both* rule sets. Participants also received feedback on each trial of the practice blocks throughout the experiment. In the second phase of the condition, participants performed the mixed block in which the rules randomly switched between trials (*Figure 1*).

**Figure 1.**
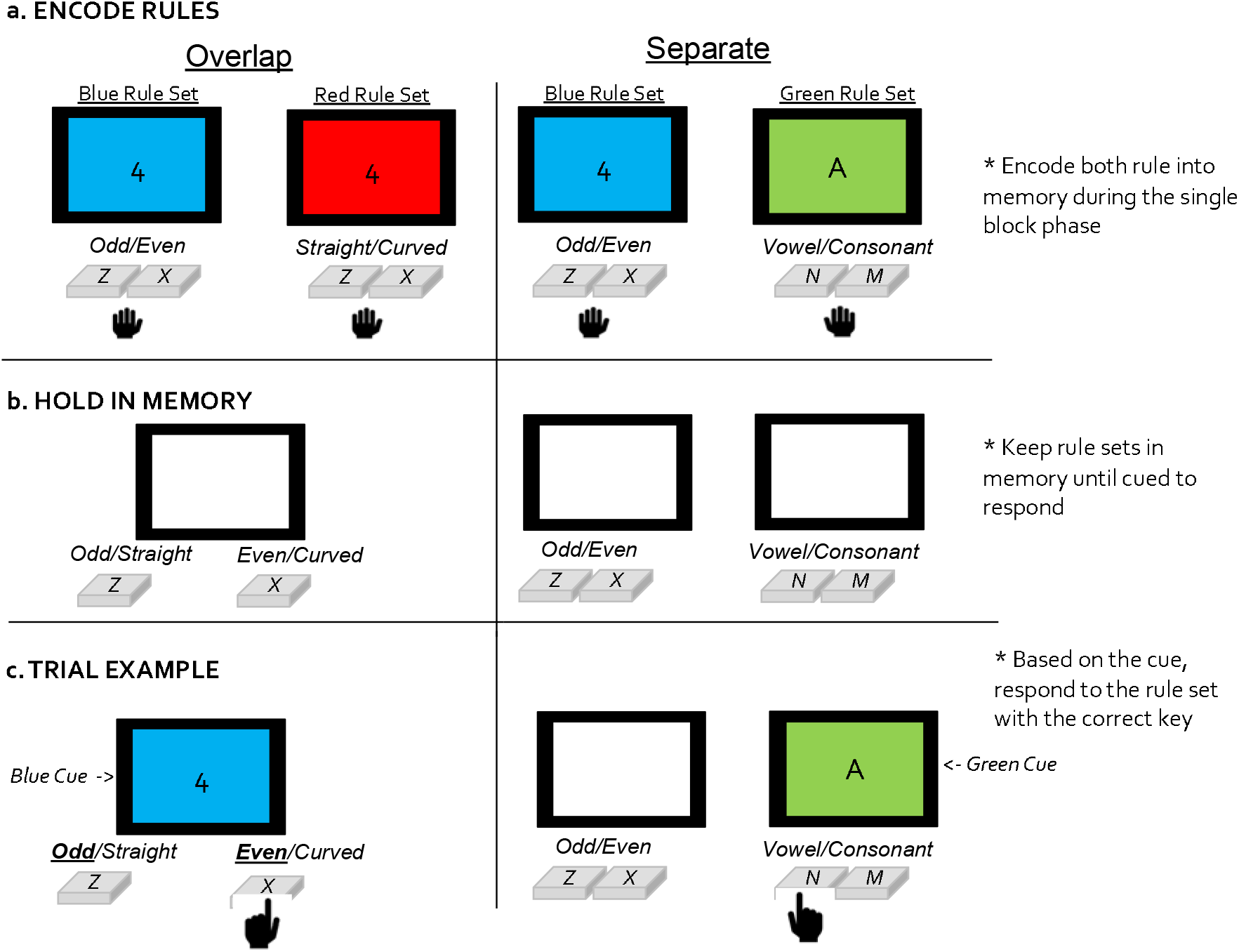
Task Switch Paradigm consists of a Separate & Overlap condition, completed on different days counterbalanced across visits within subjects. (a) Example of the Overlap condition requires a subject to make odd/even or straight/curved digit categorizations cued by blue and red respectively, while Separate condition required a digit and letter categorization of Odd/Even and Vowel/Consonant. Responses were mapped on the same hand with the middle finger corresponding to odd/straight and separately to even/curved. (b) Participants are then asked to hold both rule-sets and their responses in memory as they prepare to be randomly shown either rule-set. (c) A trial example for both conditions shows a digit stimulus paired with a blue background cue and letter stimulus paired with a green background cue which asks the participants to engage the Blue Rule Set to categorize the stimuli as Odd or Even for Overlap and in the Separate condition engage the Green Rule set to categorize the letter as Vowel or Consonant.

Preprocessing of the task-switching data consisted of validating completion of both Overlap and Separate conditions, removal of reaction times less than 200ms, and removing incorrect trials. We then calculated mixing and switching cost from this data for correct reaction times. Mixing cost was computed from performance between pure block trials and repeated trials in the mixed block. Switching cost was computed from performance between repeated and switch trials in the mixed block. These two measures were dummy coded from single, repeat, and switch trials from the task with repeated trials as the comparison trial type for single trials (e.g., mixing cost ∼ single - repeat; direction of the outputs are flipped and will be interpreted correctly to represent mixing cost) and switch trials (e.g., switching cost ∼ switch - repeat). The conditions were also dummy coded with Overlap condition serving as the base comparison for Separate condition (Overlap - Separate).

### MRI Acquisition

All MRI data were collected with a 3T GE Discovery 750W MRI scanner using a 32-channel head coil. The focus of the proposed analysis will be secondary analysis of the T2* BOLD images to characterize resting state GFC. T1 images were collected for all participants and are utilized for registration purposes. T1 images were collected via a three-dimensional fast spoiled gradient echo sequence (FSPGR) BRAVO T1 scan with the following parameters: TI=900ms, TE=3.168ms, TR=8.388ms, flip angle=8°, Acquisition Matrix=256×256×196, FOV=256×256×196, 1mm X 1mm X 1mm voxels. Participants completed three 8-minute resting state scans within each scanning session on two separate days that were at least 2 days apart. Resting state scans served as the baseline scans for an acute exercise intervention on each day. Participants were instructed not to exercise for 24 hours before their scan. Participants were instructed to attend their eyes on a fixation cross, relax, and remain awake. Resting state functional images were acquired with a voxel size of 3.44×3.44×4mm with ascending axial slice acquisition and no gap between slices (TE=30ms, TR=2000ms, flip angle= 80°, FOV=256×256mm, Matrix size=64×64×37; 240 volumes for a total scan time of 480sec). Each functional imaging run began with five dummy volumes that were removed prior to the nuisance regression. Therefore, in total, all participants had six 8-minute resting state scans across two days, for a total of 48 minutes and between 705 to 1,410 volumes (small number of subjects had runs dropped due to imaging artifacts).

### Resting state preprocessing

Resting state image processing utilized FMRIPREP v22.0.2 (Esteban et al., 2019), a Nipype based tool. Each T1-weighted volume was corrected for INU (intensity non-uniformity) using N4BiasFieldCorrection and skull-stripped using antsBrainExtraction.sh (using the Oasis30ANTs template). Brain surfaces from FreeSurfer v7.1, and the brain mask estimated previously was refined with a custom variation of the method to reconcile ANTs-derived and FreeSurfer-derived segmentations of the cortical gray-matter of Mindboggle (Klein et al., 2017). Spatial normalization to the ICBM 152 Nonlinear Asymmetrical template version 2009c (Fonov et al., 2011) was performed through nonlinear registration using ANTs (Avants et al., 2008) with brain-extracted versions of both the T1-weighted volume and the ICBM template. Brain tissue segmentation of cerebrospinal fluid (CSF), white-matter (WM) and gray-matter (GM) was performed on the brain-extracted T1-weighted images using fast (FSL v5.0.9). Volume-based spatial normalization to two standard spaces (MNI152NLin2009cAsym, MNI152NLin6Asym) was performed through nonlinear registration with antsRegistration (ANTs 2.3.3), using brain-extracted versions of both T1w reference and the T1w template.

Next, for each of the six resting state BOLD runs per subject, the following preprocessing was performed. First, a reference volume and its skull-stripped version were generated using a custom methodology of fMRIPrep. Susceptibility distortion correction (SDC) was omitted. The BOLD reference was then co-registered to the T1w reference using *bbregister* (FreeSurfer) which implements boundary-based registration (Greve & Fischl, 2009). Co-registration was configured with six degrees of freedom. Head-motion parameters with respect to the BOLD reference (transformation matrices, and six corresponding rotation and translation parameters) are estimated using mcflirt before any spatiotemporal filtering (FSL 5.0.9) (Jenkinson et al., 2002). The BOLD time-series were resampled onto their original, native space by applying the transforms to correct for head-motion.

These resampled BOLD time-series will be referred to as preprocessed BOLD in original space, or just preprocessed BOLD. The BOLD time-series were resampled into standard space, generating a preprocessed BOLD run in MNI152NLin2009cAsym space. All resamplings can be performed with a single interpolation step by composing all the pertinent transformations (i.e., head-motion transform matrices, and co-registrations to anatomical and output spaces). Gridded (volumetric) resamplings were performed using antsApplyTransforms (ANTs), configured with Lanczos interpolation to minimize the smoothing effects of other kernels (Lanczos, 1964). Many internal operations of fMRIPrep use Nilearn 0.6.2 (Abraham et al., 2014) (RRID:SCR_001362), mostly within the functional processing workflow. For more details of the pipeline, see the section corresponding to workflows in fMRIPrep’s documentation.

### Denoising for functional connectivity analyses

Additional preprocessing and data preparation for functional connectivity analyses was performed with xcpEngine (https://xcpengine.readthedocs.io/index.html), v1.2.4. (Ciric et al., 2018) [https://github.com/PennLINC/xcpEngine/blob/master/designs/fc-acompcor.dsn]. In brief, the anatomical compcor design was used, with the global signal added as an additional nuisance regressor. First, the first five volumes of each scan and corresponding nuisance regressors were removed. Temporal filtering (3dBandpass function from AFNI, butterworth filter) was done to ensure the BOLD fluctuations are within the frequency band of 0.01 < f < 0.08 Hz that has been identified as optimally sensitive to functional connectivity associated with brain networks. Nuisance regression resulted in residualized volumes which is the left-over signal after removing nuisance regressors.

### Global functional connectivity analyses

Global functional connectivity (GFC) is computed as a region’s connectivity to every other region of the brain similar to the method of Global Brain Connectivity (Cole et al., 2012). First, for every subject, Schaefer Atlas grey matter masks of the residualized volumes from the xcpEngine output were isolated from the whole brain to focus on the cortical gray matter regions of the brain. Second, using AFNI’s 3dTcorrMap, the timeseries in each voxel in grey matter was correlated with the timeseries in every other voxel (excluding itself) to create GFC voxelwise maps (Schaefer et al., 2018).

The GFC value represents the average of all corresponding pair-wise correlations. Importantly, as averaging positive and negative values would cancel each other out, in line with previous approaches we only used positive correlation values to compute the GFC value per voxel (Cole et al., 2012; Franzmeier et al., 2017). Third, the Yeo 7 network atlas was used to create a binary mask of the cognitive control network and Default Mode Network (DMN), from which GFC per voxel within these masks was used to compute GFC values per run and then averaged across six runs for each network using *fslmaths (Yeo et al., 2011)*. The DMN and SMN were used as a comparison and only negative and absolute correlation values respectively were used (Cole et al., 2012). Using output from fMRIPrep, a framewise displacement cutoff of less than 0.5 was used per resting state run.

### Brain-behavior data Analysis

Behavioral data and GFC measures were analyzed using linear mixed effects modeling in R to test subjects’ task-switching performance and moderating effect of CCN GFC. Linear mixed effects models with crossed predictors were applied for the behavioral data and then separately for each network by cost and by condition, with specific interest on the trial type (i.e., cost type) by network split by condition. Linear mixed models were chosen to account for individual differences between subjects’ behavioral performance and individuals brain network GFC as well as to account for trial level analysis of the outcome variable. To correct for multiple comparison, a Bonferroni threshold for reaching significance will be *p*-value of .025 (i.e., .05 divided by the 2 task conditions of the task switch paradigm) for the brain behavior models where Separate and Overlap task conditions where in separate models. Age, gender, years of education, and framewise displacement were entered as covariates. Subject outliers were removed whose accuracy was below chance (n=3), and those who dropped and did not complete behavioral data (n=6) with a final sample of 113.

**Table 1.**
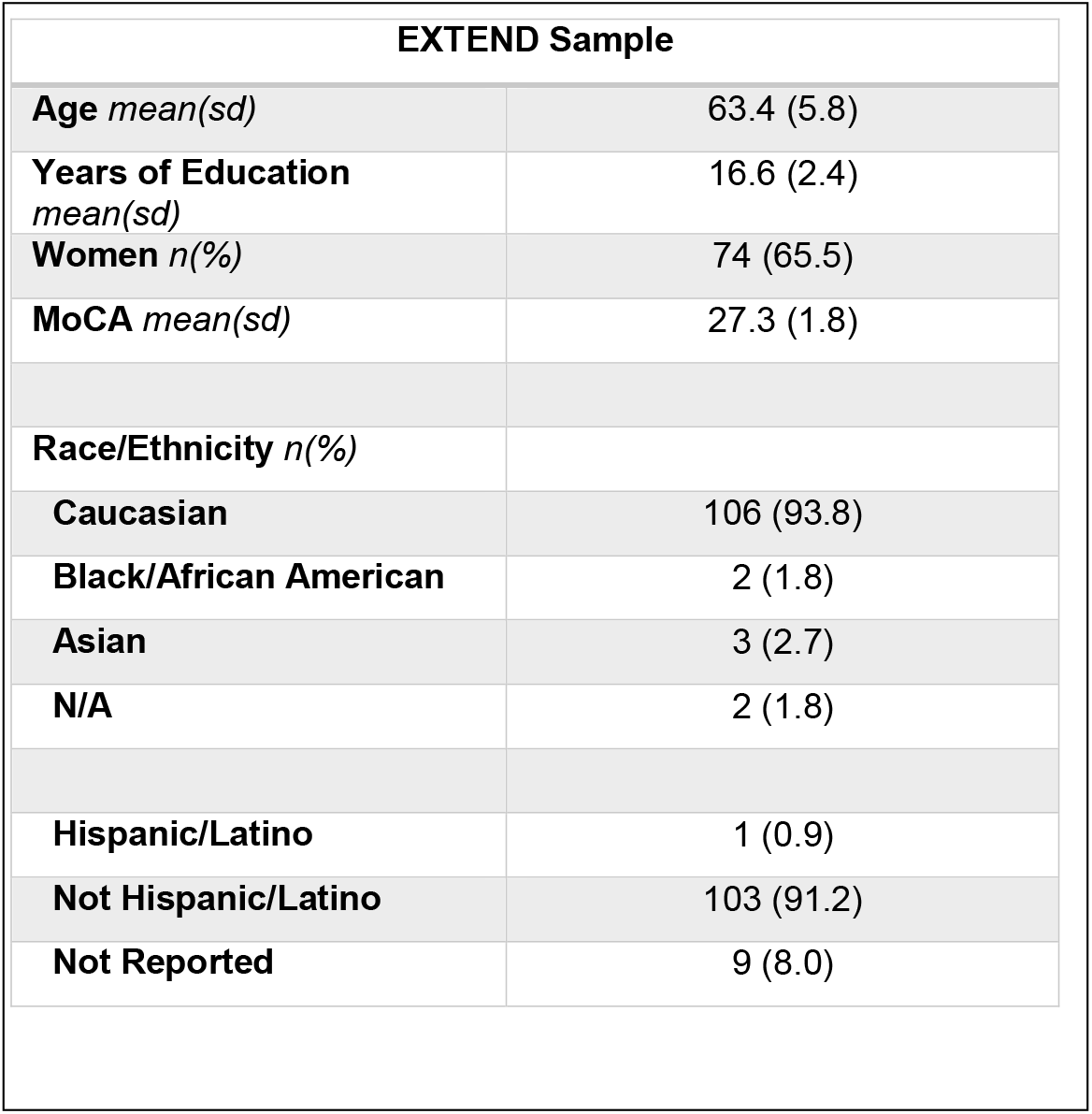
Study Demographics.

| EXTEND Sample |  |
| --- | --- |
| **Age** *mean(sd)* | 63.4 (5.8) |
| **Years of Education** *mean(sd)* | 16.6 (2.4) |
| **Women** *n(%)* | 74 (65.5) |
| **MoCA** *mean(sd)* | 27.3 (1.8) |
| **Race/Ethnicity** *n(%)* |  |
| **Caucasian** | 106 (93.8) |
| **Black/African American** | 2 (1.8) |
| **Asian** | 3 (2.7) |
| **N/A** | 2 (1.8) |
| **Hispanic/Latino** | 1 (0.9) |
| **Not Hispanic/Latino** | 103 (91.2) |
| **Not Reported** | 9 (8.0) |

## 3. Results

The results describe five main analyses. First, to understand the effect of our behavioral manipulation we examined the relationship (1) between task condition (i.e., Separate and Overlap) and trial type (i.e., single, repeated, and switch) on correct trial reaction times. Next, to understand the effect of network GFC on behavioral performance we separated our data by condition and cost to understand the relationship between GFC and cost type within each condition. The Overlap was split into (2) Mixing and (2) Switching cost models and repeated for the Separate condition. The four models tested the relationship of network GFC (CCN, DMN, SMN) on trial type (Mixing and Switching cost) on our outcome of correct RT. To address RT skewness, we applied logarithmic transformation to the RT data. For the network interactions, Bonferroni multiple comparisons was applied for the two task conditions (i.e., Separate and Overlap).

### Behavioral performance

Our first study aim was to examine the impact of overlapping response mappings by comparing Separate and Overlap conditions (Table 2). Age was a significant predictor of reaction time (β = 0.029, *t*(105.06) = 2.46, *p* = 0.016) such that older adults responded slower. Years of education and gender were not significant predictors (YrsEdu: β = −0.012, *t*(105.02) = −1.06, *p* = 0.289, gender: β = −0.015, *t*(104.96) = −0.64, *p* = 0.526) such that older adults reaction time on the task slowed independent of education and gender. Consistent with our prediction, there was a significant interaction between task condition on trial type costs, such that the overlap condition showed increased mixing cost (β = 0.238, *t*(42671.94) = 31.88, *p* < 0.001) but not switching costs (β = 0.003, *t*(42671.92) = 0.54, *p* > 0.59) compared to the separate condition (Figure 2).

**Figure 2.**
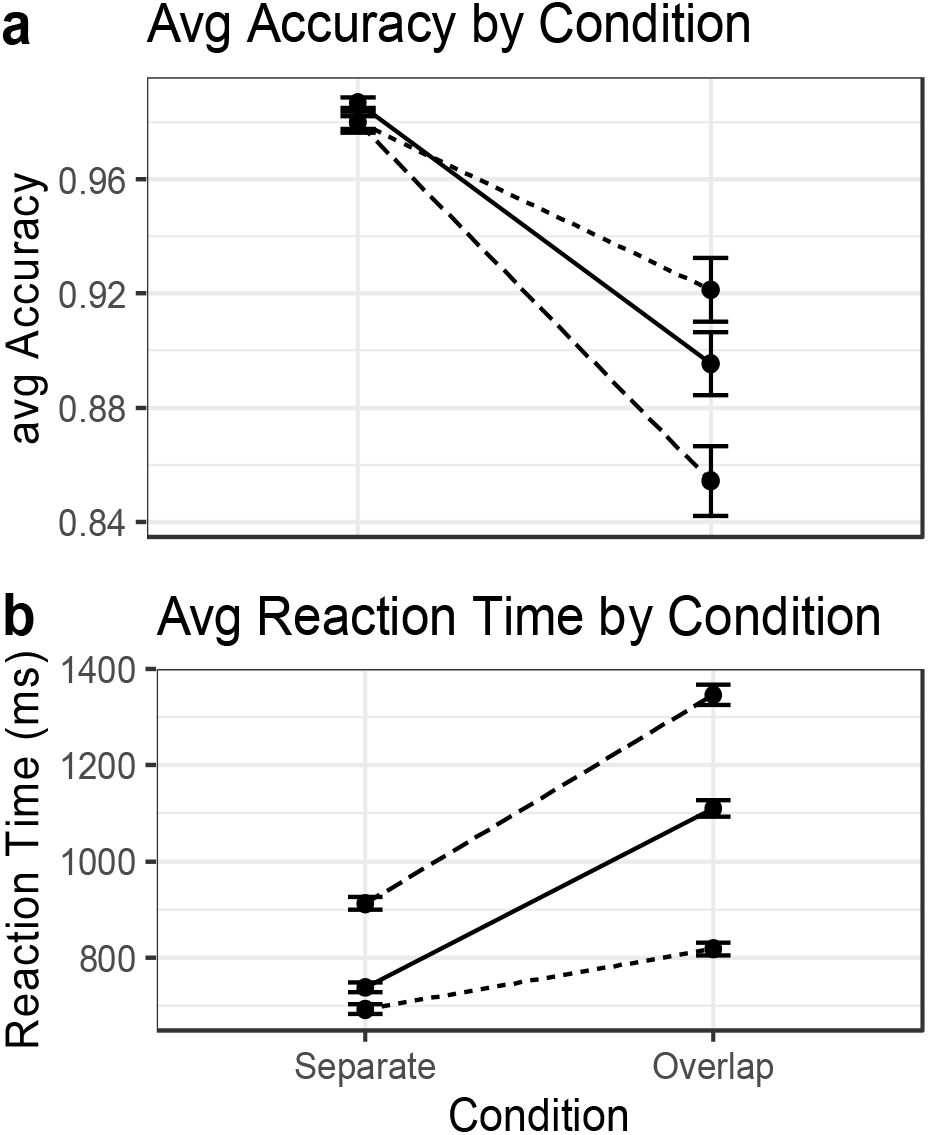
Line graph showing average mean reaction times for each trial type (Single, Repeated, and Switched) by condition (Separate and Overlap). Differences between Repeated and Single indicate Mixing cost while differences be-tween Switch and Repeated indicate Switching cost. Dotted line = Single; Solid = Repeated; Dashed = Switched Trials.

**Table 2.**
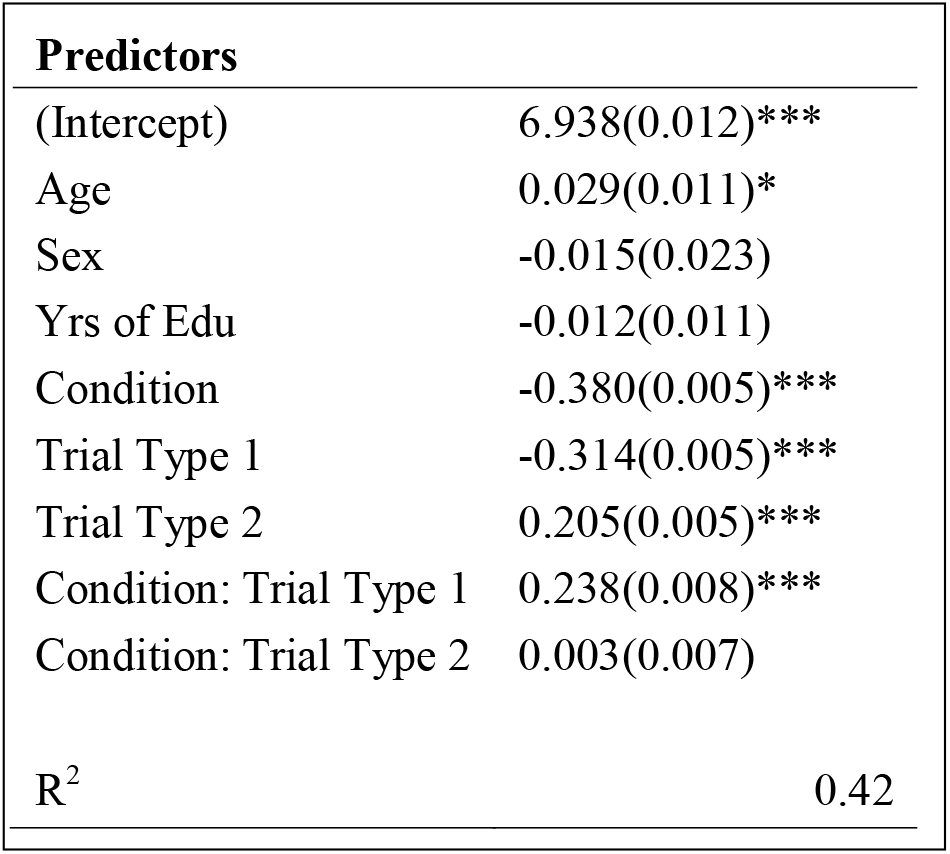
All linear mixed models include age, sex, and years of education as covariates. Age measured as chronological age in years; sex coded as M=1, F=0. Beta estimates (Std. Error). All continuous variables were mean centered for interpretability of the interactions. The table above shows linear mixed models with respect to task-switch performance for log corrected reaction time, mixing cost (calculated as Single trials – Repeat trials; Trial Type 1), and switching cost (calculated as Switch trials – Repeated trials; Trial Type 2). Note that Condition contrast was set as Separate – Overlap

### Network GFC relationship on Mixing Cost

Our primary interest for this study was to determine if CCN GFC would relate to task switching performance, predicting lower mixing cost in the overlap condition compared to separate condition.

The following covariates in the model now reflect the inclusion of framewise displacement (FD) and GFC. For the relationship between network GFC and Mixing cost in the Overlap condition (Table 3), Age was a significant predictor of reaction time (β = 0.034, *t*(106.82) = 2.61, *p* = 0.010) such that older adults responded slower. Gender and years of education were not significant predictors (Gender: β = −0.022, *t*(106.33) = −0.79, *p* = 0.427, YrsEdu: β = −0.030, t(106.71) = −2.23, *p* = 0.027) such that older adults reaction time on the task slowed independent of gender and years of education. Framewise Displacement was not significant predictor (β = −0.064, *t*(106.23) = −0.40, *p* = 0.682) such that older adults reaction time on the task slowed independent of framewise displacement. There was a significant interaction between network GFC and trial type for Mixing cost (CCN: β = −0.012, *t*(13176.93) = −2.04, *p* = 0.040, DMN: β = 0.015, *t*(13177.86) = 2.52, *p* = 0.011, SMN: β = −0.000, *t*(13176.91) = −0.11, *p* = 0.911) indicating older adults with greater CCN and SMN GFC did not show a relationship with mixing cost but greater anticorrelated DMN GFC strength did relate to better performance (Figure 3).

**Figure 3.**
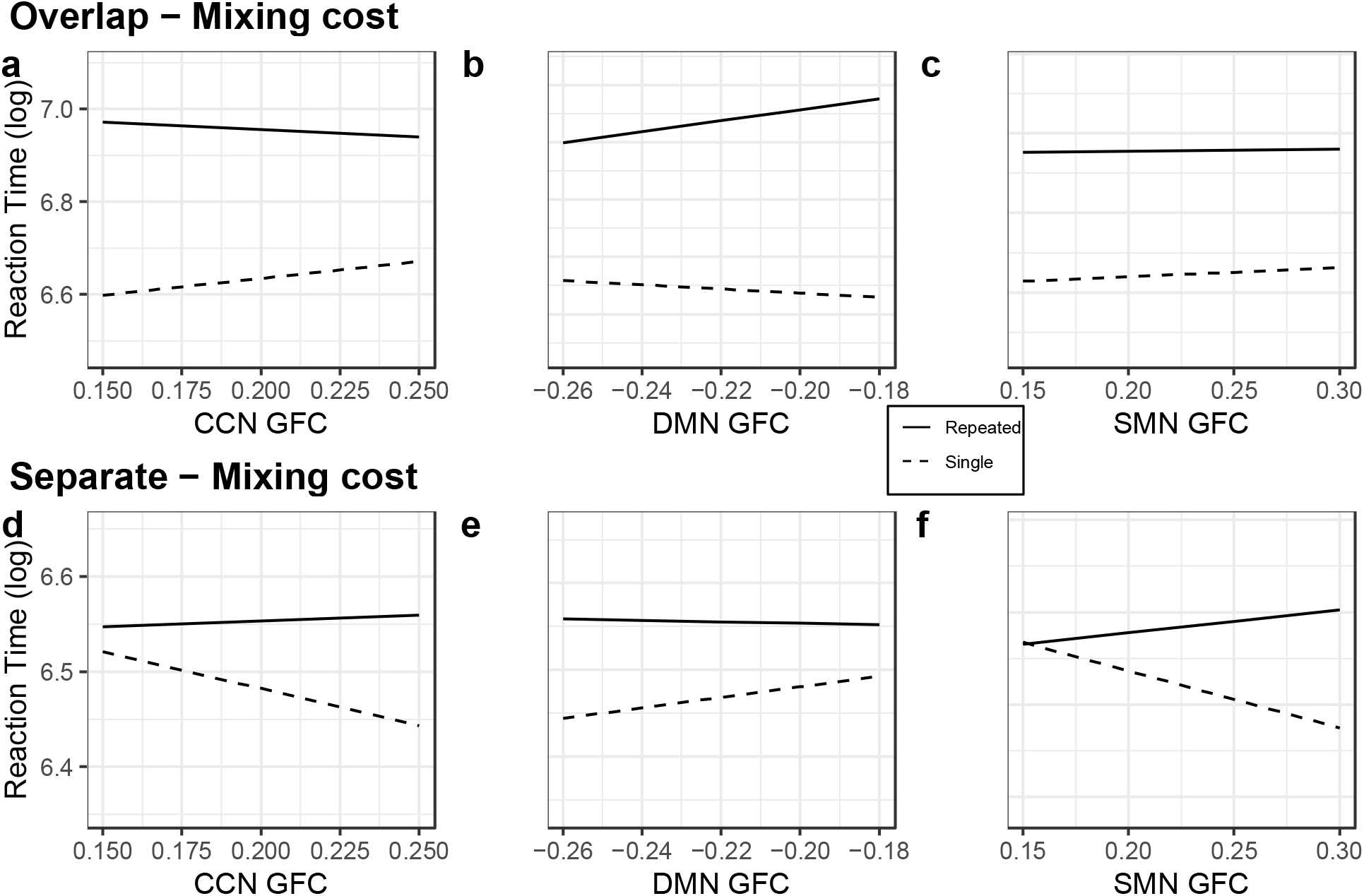
Graphs show differences between mean log transformed reaction times of Repeated (solid line) and Single (dashed line) trials, mixing cost in both conditions, with the predicted values and their confidence intervals. a) Increasing CCN GFC showed decreasing mixing cost b) decreasing DMN GFC showed decreasing mixing cost and c) non-significant increasing SMN GFC showed decreasing mixing cost. d) increasing CCN GFC showed increasing mixing cost. e) increasing negative DMN GFC showed increasing mixing cost. f) increasing SMN GFC showed increasing cost. Findings show CCN and DMN GFC relate to performance for Overlap mixing cost but not SMN GFC while in the Separate condition we see poor performance with all three networks.

**Table 3.** All linear mixed models include age, sex, and years of education as covariates. Age measured as chronological age in years; sex coded as M=1, F=0. Beta estimates (Std. Error). All continuous variables were mean centered for interpretability of the interactions. The table above shows linear mixed models with respect to task-switch performance for log corrected reaction time, mixing cost (calculated as Single trials – Repeat trials; Trial.Type,) in both Overlap and Separate condition. Betas should be interpreted in the opposite direction for Mixing Cost.

| Overlap Mixing Cost |  |  |  |
| --- | --- | --- | --- |
| Predictors | CCN GFC | DMN GFC | SMN GFC |
| (Intercept) | 6.935(0.014)*** | 6.935(0.014)*** | 6.935(0.014)*** |
| Age | 0.034(0.013)* | 0.035(0.013)** | 0.034(0.013)* |
| Sex | -0.022(0.028) | -0.021(0.028) | -0.022(0.028) |
| Yrs of Edu | -0.030(0.013)* | -0.030(0.013)* | -0.030(0.013)* |
| FD | -0.064(0.157) | -0.063(0.156) | -0.063(0.160) |
| NetGFC | -0.004(0.013) | 0.012(0.013) | -0.000(0.014) |
| Trial.Type | 0.314(0.006)*** | 0.314(0.006)*** | 0.314(0.006)*** |
| NetGFC:Trial.Type | -0.012(0.006)* | 0.015(0.006)* | -0.000(0.006) |
| R<sup>2</sup> |  |  | 0.24 |
| Separate Mixing Cost |  |  |  |
| Predictors | CCN GFC | DMN GFC | SMN GFC |
| (Intercept) | 6.561(0.012)*** | 6.560(0.012)*** | 6.561(0.012)*** |
| Age | 0.034(0.011)** | 0.034(0.011)** | 0.034(0.11)** |
| Sex | 0.003(0.025) | 0.002(0.025) | 0.003(0.025) |
| Yrs of Edu | -0.002(0.012) | -0.002(0.012) | -0.002(0.012) |
| FD | 0.126(0.139) | 0.121(0.138) | 0.131(0.141) |
| NetGFC | -0.000(0.011) | 0.000(0.011) | 0.004(0.012) |
| Trial.Type | -0.076(0.004)*** | -0.076(0.004)*** | -0.076(0.004)*** |
| NetGFC:Trial.Type | 0.009(0.004)* | -0.008(0.004)* | 0.024(0.004)*** |
| R<sup>2</sup> |  |  | 0.21 |
| *** p < 0.001; ** p < 0.01; * p < 0.05; . p < 0.10 |  |  |  |

For the relationship between network GFC and Mixing cost in the Separate condition, Age was a significant predictor of reaction time (β = 0.034, *t*(106.95) = 2.85, *p* = 0.005) such that older adults responded slower. Gender and Years of education were not significant predictors (Gender: β = 0.003, *t*(106.94) = 0.12, *p* = 0.901, YrsEdu: β = −0.002, *t*(106.98) = −0.22, *p* = 0.819) such that older adults reaction time on the task slowed independent of Gender and Education. Framewise Displacement was not significant predictor (β = 0.126, *t*(106.82) = 0.91, *p* = 0.364) such that older adults reaction time on the task slowed independent of framewise displacement. There was a significant interaction between network GFC on Mixing cost (CCN: β = 0.009, *t*(14296.29) = 2.20, *p* = 0.027, DMN: β = −0.008, *t*(14296.32) = −2.04, *p* = 0.041, SMN: β = 0.024, *t*(14296.67) = 5.53, *p* < 0.001) indicating older adults with greater SMN GFC showed poorer performance (i.e., greater mixing cost) while CCN and DMN showed similar pattern but did not meet the statistical threshold of significance.

These results show that in our more challenging condition in which response rule set boundaries are overlapped, older adults show a different pattern of GFC for mixing cost depending on condition difficulty. For the Overlap condition, the relationship between DMN GFC emerges with better performance but not CCN and SMN. This finding falls in line with prior work showing Mixing cost cannot be explained by processing speed alone which could be most related to SMN though surprisingly CCN, a network related to EF abilities, does not show a significant relationship with performance. In the Separate condition, a relationship between SMN emerges with poorer performance. DMN and CCN GFC does not reach the multiple comparisons threshold for significance with mixing cost in the Separate condition. This finding may be combining two factors, the complex cognitive mechanism needed for mixing cost (DMN) and the increase in rule set key responses from two (Overlap) to four keys in the Separate condition maybe decreasing cognitive demands as there is less interference being created at the rule-set and key response sets.

**Figure 4.**
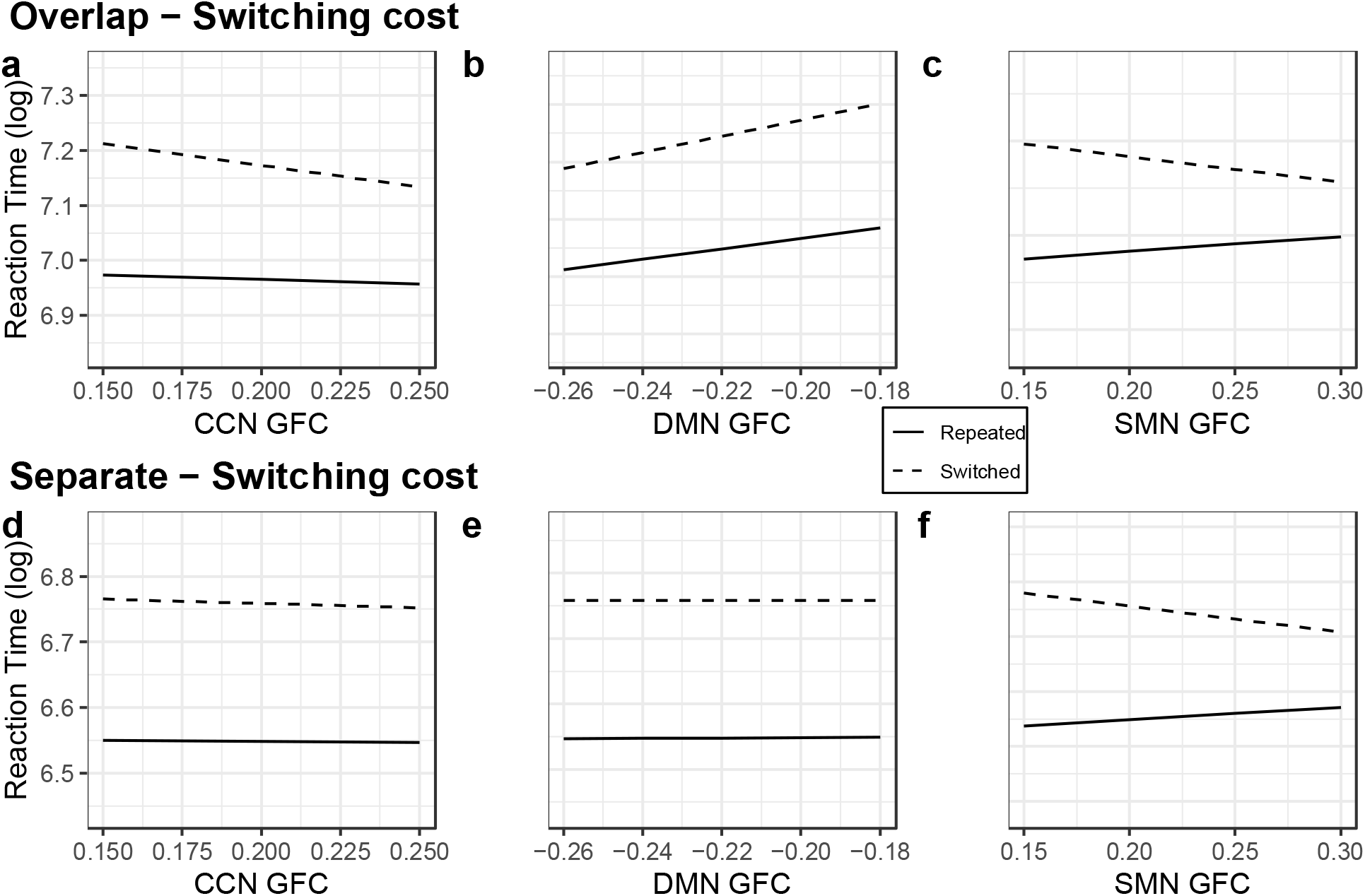
Graphs show differences between log transformed reaction times of Repeated (solid line) and Switch (dashed line) trials, Switching cost, with the predicted values and their confidence intervals. a) Non-significant increasing CCN GFC showed no effect on switching cost b) decreasing DMN GFC showed no effect switching cost and c) increasing SMN GFC showed a significant decrease in Switching cost. Findings show a non-significant relationship between CCN and DMN GFC to performance but SMN GFC did relate to better performance in Separate switch cost.

**Table 4.**
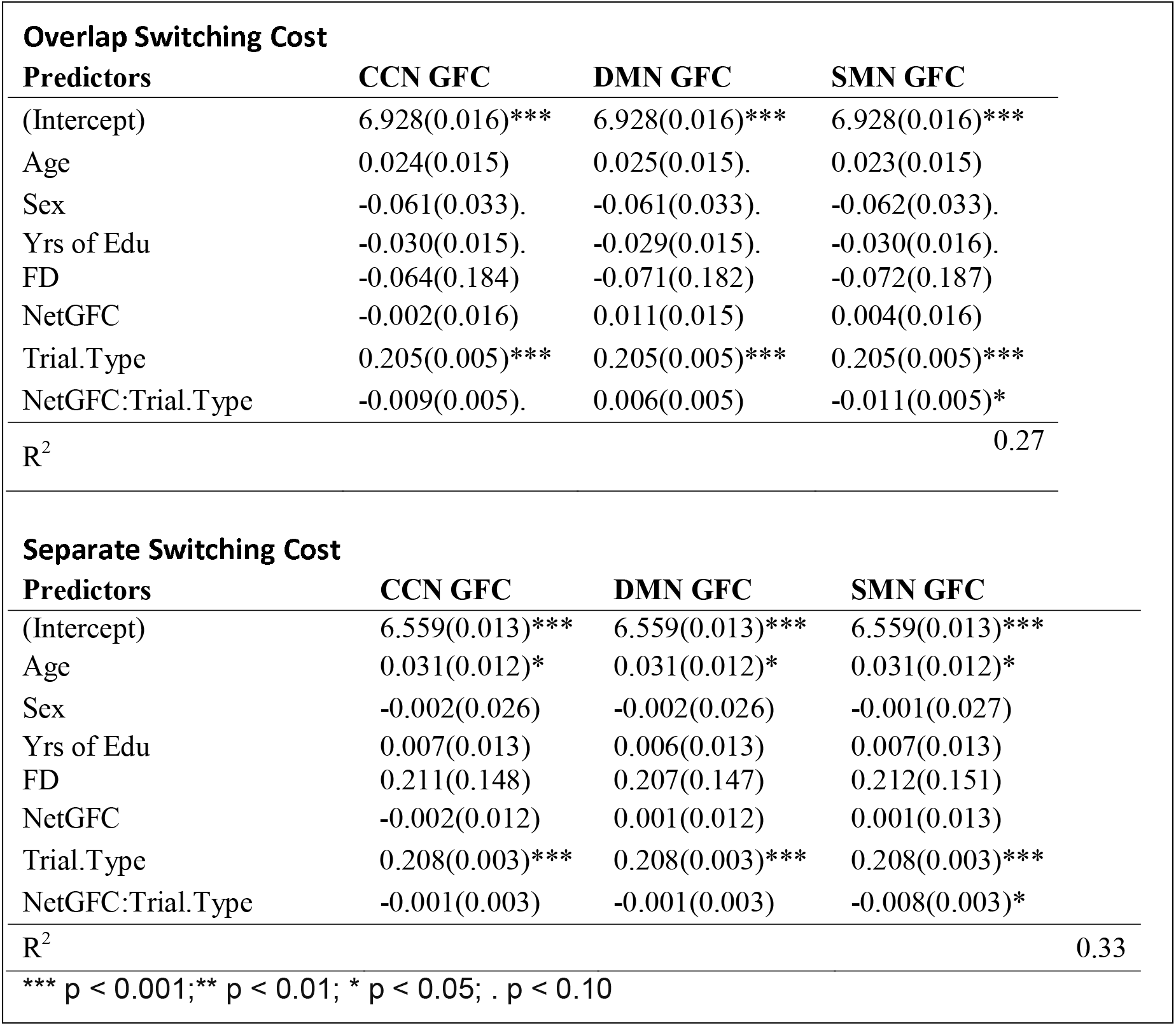
All linear mixed models include age, sex, and years of education as covariates. Age measured as chronological age in years; sex coded as M=1, F=0. Beta estimates (Std. Error). All continuous variables were mean centered for interpretability of the interactions. The table above shows linear mixed models with respect to task-switch performance for log corrected reaction time, switching cost (calculated as Switch trials – Repeated trials; Trial.Type).

### Network GFC relationship on Switching Cost

Compared to our primary question in the prior section, we predicted that CCN GFC would not be related to lower switching cost in the Overlap or Separate condition.

For the relationship between network GFC and switching cost in Overlap condition, Age, Years of education, and gender were not significant predictors of reaction time (Age: β = 0.024, *t*(106.90) = 1.60, *p* = 0.112, YrsEdu: β = −0.030, *t*(106.74) = −1.88, *p* = 0.061, Gender: (β = −0.061, *t*(106.35) = −1.85, *p* = 0.066) such that reaction time on the task were independent of Age, Education, and Gender. Framewise Displacement was not significant predictor (β = −0.064, *t*(106.31) = −0.34, *p* = 0.727) such that older adults reaction time on the task slowed independent of framewise displacement. There was a significant interaction between network GFC on Switching cost (CCN: β = −0.009, *t*(15799.69) = −1.77, *p* = 0.076, DMN: β = 0.006, *t*(15799.46) = 1.19, *p* = 0.231, SMN: β = −0.011, *t*(15802.01) = −2.24, *p* = 0.024) indicating greater SMN GFC showed lower Switching cost (i.e., better performance) but not with CCN or DMN GFC.

For the relationship between network GFC and switching cost in Separate condition, Age was a significant predictor of reaction time (Age: β = 0.031, *t*(107.02) = 2.47, *p* = 0.014) such that older adults responded slower. Gender and Years of education were not significant predictors (Gender: β = 0.002, *t*(106.99) = 0.07, *p* = 0.937, YrsEdu: β = 0.007, *t*(107.01) = 0.53, *p* = 0.590) such that older adults reaction time on the task slowed independent of Gender, and Education. Framewise Displacement was not significant predictor (β = 0.211, *t*(106.98) = 1.41, *p* = 0.158) such that older adults reaction time on the task slowed independent of framewise displacement. There was a significant interaction between network GFC on Switching cost (CCN: β = −0.001, *t*(17842.89) = −0.29, *p* = 0.767, DMN: β = −0.001, *t*(17843.10) = −0.35, *p* = 0.719, SMN: β = −0.008, *t*(17845.83) = −2.38, *p* = 0.017) indicating older adults with greater SMN GFC showed lower switching cost (i.e., better performance) but not with CCN or DMN GFC.

These results show SMN GFC relationship on switching cost in the Overlap and Separate condition predicting better performance. The SMN GFC may be attributed to an individual’s processing speed whether it be in a condition with response rule set boundaries are overlapped or clearly delineated. In the Overlap condition CCN and DMN did not relate to better performance for switching cost as it did not reach significance while in the Separate condition the relationship was much weaker.

## 4. Discussion

Based on previous work showing CCN’s GFC relating to higher cognitive control abilities, we aimed to determine how age-related EF performance would relate to differences in resting state FC of CCN measured by GFC. Primarily, we predicted that in order to perform well in the cognitive demanding condition of a task switch paradigm, individuals with greater CCN GFC would perform better. We found greater anticorrelated strength of DMN GFC (i.e., negative values) predicted lower mixing cost in the overlap condition. We also found that greater SMN GFC predicted worse performance, or greater mixing cost, in the separate condition. Only the relationship between greater SMN GFC and switching cost predicted better performance in both conditions.

### 4.1. Age-related Differences in network GFC and Mixing Cost

We observed selective relationships between mixing cost and network GFC in which stronger anticorrelated DMN GFC related to better performance in the Overlap condition but not in the Separate condition. Contrary to our prediction CCN GFC did not meet the multiple comparison statistical threshold for significance in the Overlap condition and did not meet the threshold for the trend to worse performance in the Separate condition. The SMN GFC relationship was related to poorer performance in the Separate condition. Our task switch paradigm replicated the effects of overlapping response-rule set as shown by Mayr (2001). Research on task-switching, aging (Mayr, 2001; Salthouse et al., 1998), and the relationship between CCN GFC and cognitive control performance (Cole et al., 2012; Franzmeier et al., 2017), led us to predict that greater CCN GFC would predict lower mixing cost performance in the Overlap condition compared to the Separate condition. We did not find CCN GFC to be related to better performance in our cohort of healthy older adults and instead it was DMN GFC that was related to better performance for mixing cost in the Overlap condition.

### 4.2. Age-related Differences in network GFC and Switching Cost

For switching cost, while not meeting our multiple comparison threshold, greater SMN GFC was related to better performance in both conditions which alludes to prior studies identifying this measure to be related to processing speed (Wasylyshyn et al., 2011). Switching cost has been shown to be related more so to the speed of switching from one rule set to another and therefore acting upon the desired rule in the moment as quickly as possible requiring the SMN to engage the finger press. Greater CCN GFC and the anticorrelated strength of DMN GFC was not related to performance in either condition.

### 4.3. Task Switching cognitive processes and Network GFC

The performance decline of mixing cost was shown to be mitigated by GFC of the anticorrelated strength of the DMN. Previous studies have shown mixing cost decreases due to age-related deficits above and beyond slowing of processing speed. Mitigation of mixing cost decline may be due DMN’s role in complex higher order EF processes needed to perform the task which involves working memory (Kelly et al., 2008; Wasylyshyn et al., 2011; Zou et al., 2013). Furthermore, the relationship of DMN GFC with mixing cost was not seen with switching cost where we know declines are closer related to processing speed and less on age-related deficits of higher-order processes. The increased difficulty in switching between tasks when rule-response mappings are ambiguous, compared to when they are distinct, may be taxing working memory and episodic binding which have been shown to also be related to the DMN. This is particularly true in the Overlap condition, where individuals are required to hold two tasks in mind and map them onto the same effectors (i.e., fingers). Additionally, in the Overlap condition, individuals must keep track of the two conflicting tasks on the same finger, even when inhibiting the irrelevant task.

In support of switching cost and its relationship to general slowing, we found that there was selective relationship trend for greater SMN GFC in both Overlap and Separate conditions predicting lower switching cost performance. SMN functions related to motor control and physical actions may have influenced the speed at which individuals could quickly and accurately perform the task rather than affecting the higher order processes that underlie mixing cost (i.e., EF)(Biswal et al., 1995).

It is also important to note the strength of this task and how it lent itself to be leveraged in this study. Our behavioral results were consistent with previous findings in which declines in switching cost have been observed to be related to processing speed ability and mixing cost to age-related deficits above and beyond declines in speed related to more complex higher order functions. While we saw that switching cost increased by condition, the difference between the trial types to calculate switching cost (i.e., Switch - Repeat) remained similar across both conditions. Mixing cost however showed a difference between repeated and single trials from Separate to Overlap condition. This falls in line with the theory that switching cost has an additive effect to decline while mixing cost has a multiplicative effect to decline (Verhaeghen, 2011). Another feature that supported our novel results is the strength of within-subject measures of both conditions. In the measures we expected age related declines behaviorally, we can infer that performance for our sample in the Separate condition was truncated making it difficult to clearly identify high performers from low performers. The increased difficulty of the Overlap condition allowed us to see performance stratify allowing us to clearly see high performers from low performers and in turn identify relationships with better preserved brain function as measured by GFC. Collectively, these results show the nature by which task manipulation allows us to view which processes affected by aging are related to preserved brain connectivity in older adults and indicates resting state DMN GFC as a brain feature for cognitive ability in aging.

### 4.4. Limitations

We are unable to make strong aging effect claims as we did not have a cross sectional study with young adults. While we can infer based on previous cross-sectional studies, in our current study with only older adults we can only speak to the strong manipulations of the task and its effects on our sample. A strength in our study is that we had within-subject repeated measures and counter balanced our conditions, however, we are unable to account for crossover effects. Even with counter balancing the conditions there may be different effects across individuals that are not controllable in our current study. Additionally, we must be aware that our claims would be stronger if we had both resting state and task activation given that our resting state information gives us a baseline connectivity signal not activated by task, as well as uncertainty about an individual’s mental state in scanner that could influence resting state network strengths. With both rest and task state imaging data we could confidently make inferences regarding executive function having a relation with greater network GFC and less task activation relating to better performance.

### 4.5. Conclusions

This study’s findings inform our understanding of EF, and cognitive control broadly, in aging by determining DMN’s functional involvement globally to carry out higher order cognitive processes by leveraging our task switch paradigm. By using a task switch-paradigm with two conditions with increasing difficulty and GFC to measure the global strength of networks associated with the task we have shown the ability to relate rs-fc to EF ability in healthy older adults. Additional work is needed however to fully understand the mechanism at play in aging populations. Future work should look to implement both task and rest to show the relationship between those two states. Further understanding network specific characteristics and its relationship to behavioral tasks is needed and may benefit from using modulatory analysis to probe if performance is better predicted by within or between network connectivity during this aging period.

## Bibliography & References Cited

Abraham, A., Pedregosa, F., Eickenberg, M., Gervais, P., Mueller, A., Kossaifi, J., Gramfort, A., Thirion, B., & Varoquaux, G. (2014). Machine learning for neuroimaging with scikit-learn. Front Neuroinform, 8, 14. 10.3389/fninf.2014.00014

Andrews-Hanna, J. R., Smallwood, J., & Spreng, R. N. (2014). The default network and self-generated thought: component processes, dynamic control, and clinical relevance. Ann N Y Acad Sci, 1316(1), 29–52. 10.1111/nyas.12360

Avants, B. B., Epstein, C. L., Grossman, M., & Gee, J. C. (2008). Symmetric diffeomorphic image registration with cross-correlation: evaluating automated labeling of elderly and neurodegenerative brain. Med Image Anal, 12(1), 26–41. 10.1016/j.media.2007.06.004

Badre, D. (2008). Cognitive control, hierarchy, and the rostro-caudal organization of the frontal lobes. Trends Cogn Sci, 12(5), 193–200. 10.1016/j.tics.2008.02.004

Biswal, B., Yetkin, F. Z., Haughton, V. M., & Hyde, J. S. (1995). Functional connectivity in the motor cortex of resting human brain using echo-planar MRI. Magn Reson Med, 34(4), 537–541. 10.1002/mrm.1910340409

Breukelaar, I. A., Griffiths, K. R., Harris, A., Foster, S. L., Williams, L. M., & Korgaonkar, M. S. (2020). Intrinsic functional connectivity of the default mode and cognitive control networks relate to change in behavioral performance over two years. Cortex, 132, 180–190. 10.1016/j.cortex.2020.08.014

Chan, M. Y., Park, D. C., Savalia, N. K., Petersen, S. E., & Wig, G. S. (2014). Decreased segregation of brain systems across the healthy adult lifespan. Proc Natl Acad Sci U S A, 111(46), E4997–5006. 10.1073/pnas.1415122111

Ciric, R., Rosen, A. F. G., Erus, G., Cieslak, M., Adebimpe, A., Cook, P. A., Bassett, D. S., Davatzikos, C., Wolf, D. H., & Satterthwaite, T. D. (2018). Mitigating head motion artifact in functional connectivity MRI. Nat Protoc, 13(12), 2801–2826. 10.1038/s41596-018-0065-y

Cole, M. W., Anticevic, A., Repovs, G., & Barch, D. (2011). Variable global dysconnectivity and individual differences in schizophrenia. Biol Psychiatry, 70(1), 43–50. 10.1016/j.biopsych.2011.02.010

Cole, M. W., Bassett, D. S., Power, J. D., Braver, T. S., & Petersen, S. E. (2014). Intrinsic and task-evoked network architectures of the human brain. Neuron, 83(1), 238–251. 10.1016/j.neuron.2014.05.014

Cole, M. W., Pathak, S., & Schneider, W. (2010). Identifying the brain’s most globally connected regions. Neuroimage, 49(4), 3132–3148. 10.1016/j.neuroimage.2009.11.001

Cole, M. W., & Schneider, W. (2007). The cognitive control network: Integrated cortical regions with dissociable functions. Neuroimage, 37(1), 343–360. 10.1016/j.neuroimage.2007.03.071

Cole, M. W., Yarkoni, T., Repovs, G., Anticevic, A., & Braver, T. S. (2012). Global connectivity of prefrontal cortex predicts cognitive control and intelligence. J Neurosci, 32(26), 8988–8999. 10.1523/JNEUROSCI.0536-12.2012

Dickerson, B. C., Fenstermacher, E., Salat, D. H., Wolk, D. A., Maguire, R. P., Desikan, R., Pacheco, J., Quinn, B. T., Van der Kouwe, A., Greve, D. N., Blacker, D., Albert, M. S., Killiany, R. J., & Fischl, B. (2008). Detection of cortical thickness correlates of cognitive performance: Reliability across MRI scan sessions, scanners, and field strengths. Neuroimage, 39(1), 10–18. 10.1016/j.neuroimage.2007.08.042

Eich, T. S., Rakitin, B. C., & Stern, Y. (2016). Response-Conflict Moderates the Cognitive Control of Episodic and Contextual Load in Older Adults. J Gerontol B Psychol Sci Soc Sci, 71(6), 995–1003. 10.1093/geronb/gbv046

Esteban, O., Markiewicz, C. J., Blair, R. W., Moodie, C. A., Isik, A. I., Erramuzpe, A., Kent, J. D., Goncalves, M., DuPre, E., Snyder, M., Oya, H., Ghosh, S. S., Wright, J., Durnez, J., Poldrack, R. A., & Gorgolewski, K. J. (2019). fMRIPrep: a robust preprocessing pipeline for functional MRI. Nat Methods, 16(1), 111–116. 10.1038/s41592-018-0235-4

Fonov, V., Evans, A. C., Botteron, K., Almli, C. R., McKinstry, R. C., Collins, D. L., & Brain Development Cooperative, G. (2011). Unbiased average age-appropriate atlases for pediatric studies. Neuroimage, 54(1), 313–327. 10.1016/j.neuroimage.2010.07.033

Franzmeier, N., Caballero, M. A. A., Taylor, A. N. W., Simon-Vermot, L., Buerger, K., Ertl-Wagner, B., Mueller, C., Catak, C., Janowitz, D., Baykara, E., Gesierich, B., Duering, M., Ewers, M., & Alzheimer’s Disease Neuroimaging, I. (2017). Resting-state global functional connectivity as a biomarker of cognitive reserve in mild cognitive impairment. Brain Imaging Behav, 11(2), 368–382. 10.1007/s11682-016-9599-1

Friedman, N. P., & Robbins, T. W. (2022). The role of prefrontal cortex in cognitive control and executive function. Neuropsychopharmacology, 47(1), 72–89. 10.1038/s41386-021-01132-0

Grambaite, R., Selnes, P., Reinvang, I., Aarsland, D., Hessen, E., Gjerstad, L., & Fladby, T. (2011). Executive dysfunction in mild cognitive impairment is associated with changes in frontal and cingulate white matter tracts. J Alzheimers Dis, 27(2), 453–462. 10.3233/JAD-2011-110290

Greve, D. N., & Fischl, B. (2009). Accurate and robust brain image alignment using boundary-based registration. Neuroimage, 48(1), 63–72. 10.1016/j.neuroimage.2009.06.060

Jenkinson, M., Bannister, P., Brady, M., & Smith, S. (2002). Improved Optimization for the Robust and Accurate Linear Registration and Motion Correction of Brain Images. Neuroimage, 17(2), 825–841. 10.1006/nimg.2002.1132

Kelly, A. M., Uddin, L. Q., Biswal, B. B., Castellanos, F. X., & Milham, M. P. (2008). Competition between functional brain networks mediates behavioral variability. Neuroimage, 39(1), 527–537. 10.1016/j.neuroimage.2007.08.008

Klein, A., Ghosh, S. S., Bao, F. S., Giard, J., Hame, Y., Stavsky, E., Lee, N., Rossa, B., Reuter, M., Chaibub Neto, E., & Keshavan, A. (2017). Mindboggling morphometry of human brains. PLoS Comput Biol, 13(2), e1005350. 10.1371/journal.pcbi.1005350

Lacreuse, A., Raz, N., Schmidtke, D., Hopkins, W. D., & Herndon, J. G. (2020). Age-related decline in executive function as a hallmark of cognitive ageing in primates: an overview of cognitive and neurobiological studies. Philos Trans R Soc Lond B Biol Sci, 375(1811), 20190618. 10.1098/rstb.2019.0618

Lanczos, C. (1964). A Precision Approximation of the Gamma Function. Journal of the Society for Industrial and Applied Mathematics Series B Numerical Analysis, 1(1), 86–96. 10.1137/0701008

Mackie, M. A., Van Dam, N. T., & Fan, J. (2013). Cognitive control and attentional functions. Brain Cogn, 82(3), 301–312. 10.1016/j.bandc.2013.05.004

Mayr, U. (2001). Age differences in the selection of mental sets: the role of inhibition, stimulus ambiguity, and response-set overlap. Psychol Aging, 16(1), 96–109. 10.1037/0882-7974.16.1.96

Menon, V., & D’Esposito, M. (2022). The role of PFC networks in cognitive control and executive function. Neuropsychopharmacology, 47(1), 90–103. 10.1038/s41386-021-01152-w

Nasreddine, Z. S., Phillips, N. A., Bedirian, V., Charbonneau, S., Whitehead, V., Collin, I., Cummings, J. L., & Chertkow, H. (2005). The Montreal Cognitive Assessment, MoCA: a brief screening tool for mild cognitive impairment. J Am Geriatr Soc, 53(4), 695–699. 10.1111/j.1532-5415.2005.53221.x

Rogers, R. D., & Monsell, S. (1995). Costs of a predictible switch between simple cognitive tasks. Journal of Experimental Psychology: General, 124(2), 207–231. 10.1037/0096-3445.124.2.207

Rubinov, M., & Sporns, O. (2011). Weight-conserving characterization of complex functional brain networks. Neuroimage, 56(4), 2068–2079. 10.1016/j.neuroimage.2011.03.069

Salthouse, T. A., Fristoe, N., McGuthry, K. E., & Hambrick, D. Z. (1998). Relation of task switching to speed, age, and fluid intelligence. Psychol Aging, 13(3), 445–461. 10.1037/0882-7974.13.3.445

Schaefer, A., Kong, R., Gordon, E. M., Laumann, T. O., Zuo, X. N., Holmes, A. J., Eickhoff, S. B., & Yeo, B. T. T. (2018). Local-Global Parcellation of the Human Cerebral Cortex from Intrinsic Functional Connectivity MRI. Cereb Cortex, 28(9), 3095–3114. 10.1093/cercor/bhx179

Sridharan, D., Levitin, D. J., & Menon, V. (2008). A critical role for the right fronto-insular cortex in switching between central-executive and default-mode networks. Proc Natl Acad Sci U S A, 105(34), 12569–12574. 10.1073/pnas.0800005105

Tucker-Drob, E. M. (2019). Cognitive Aging and Dementia: A Life Span Perspective. Annu Rev Dev Psychol, 1, 177–196. 10.1146/annurev-devpsych-121318-085204

Verhaeghen, P. (2011). Aging and Executive Control: Reports of a Demise Greatly Exaggerated. Curr Dir Psychol Sci, 20(3), 174–180. 10.1177/0963721411408772

Voss, M. W., Weng, T. B., Burzynska, A. Z., Wong, C. N., Cooke, G. E., Clark, R., Fanning, J., Awick, E., Gothe, N. P., Olson, E. A., McAuley, E., & Kramer, A. F. (2016). Fitness, but not physical activity, is related to functional integrity of brain networks associated with aging. Neuroimage, 131, 113–125. 10.1016/j.neuroimage.2015.10.044

Wasylyshyn, C., Verhaeghen, P., & Sliwinski, M. J. (2011). Aging and task switching: a meta-analysis. Psychol Aging, 26(1), 15–20. 10.1037/a0020912

Yeo, B. T., Krienen, F. M., Sepulcre, J., Sabuncu, M. R., Lashkari, D., Hollinshead, M., Roffman, J. L., Smoller, J. W., Zollei, L., Polimeni, J. R., Fischl, B., Liu, H., & Buckner, R. L. (2011). The organization of the human cerebral cortex estimated by intrinsic functional connectivity. J Neurophysiol, 106(3), 1125–1165. 10.1152/jn.00338.2011

Zou, Q., Ross, T. J., Gu, H., Geng, X., Zuo, X. N., Hong, L. E., Gao, J. H., Stein, E. A., Zang, Y. F., & Yang, Y. (2013). Intrinsic resting-state activity predicts working memory brain activation and behavioral performance. Hum Brain Mapp, 34(12), 3204–3215. 10.1002/hbm.22136

